# Creative tempo: Spatiotemporal dynamics of the default mode network in improvisational musicians

**DOI:** 10.1101/2024.04.07.588391

**Authors:** Harrison Watters, Abia Fazili, Lauren Daley, Alex Belden, TJ LaGrow, Taylor Bolt, Psyche Loui, Shella Keilholz

## Abstract

The intrinsic dynamics of human brain activity display a recurring pattern of anti-correlated activity between the default mode network (DMN), associated with internal processing and mentation, and task positive regions, associated with externally directed attention. In human functional magnetic resonance imaging (fMRI) data, this anti-correlated pattern is detectable on the infraslow timescale (<0.1 Hz) as a quasi-periodic pattern (QPP). While the DMN is implicated in creativity and musicality in traditional time-averaged functional connectivity studies, no one has yet explored how creative training may alter dynamic spatiotemporal patterns involving the DMN such as QPPs. In the present study, we compare the outputs of two QPP detection approaches, sliding window algorithm and complex principal components analysis (cPCA). We apply both methods to an existing dataset of musicians captured with resting state fMRI, grouped as either classical, improvisational, or minimally trained non-musicians. The original time-averaged functional connectivity (FC) analysis of this dataset used improvisation as a proxy for creative thinking and found that the DMN and visual networks (VIS) display higher connectivity in improvisational musicians. We expand upon this dataset’s original study and find that QPP analysis detects convergent results at the group level with both methods. In improvisational musicians, dynamic functional correlation in the group-averaged QPP was found to be increased between the DMN-VIS and DMN-FPN for both the QPP algorithm and complex principal components analysis (cPCA) methods. Additionally, we found an unexpected increase in FC in the group-averaged QPP between the dorsal attention network and amygdala in improvisational musicians; this result was not reported in the original seed-based study of this dataset. The current study represents a novel application of two dynamic FC detection methods with results that replicate and expand upon previous seed-based FC findings. The results show the robustness of both the QPP phenomenon and its detection methods. This study also demonstrates the value of dynamic FC methods in reproducing seed-based findings and their promise in detecting group-wise or individual differences that may be missed by traditional seed-based resting state fMRI studies.

## 1. Introduction

Intrinsic brain activity is dominated by alternating activity between brain regions associated with internal and external attention (Fox et al., 2005. Raichle, 2015a. Raichle, 2015b. Abbas et al., 2019a). The primary network known to increase activity in the absence of externally directed attention is the default mode network (DMN), which is typically considered to include the posterior cingulate cortex (PCC), medial prefrontal cortex (mPFC), and precuneus (Raichle, 2015. Smallwood et al., 2021). The DMN shows increased activity during internally directed processes such as memory replay, mind wandering, imagination of future scenarios, social inference, and the construction of shared and individual internal narratives (Zadbood et al., 2017. Buckner et al., 2019. Yeshurun., 2021. Menon et al., 2023). In global-signal regressed data from resting state functional magnetic resonance imaging (rsfMRI), activity in default mode regions is strongly anti-correlated with regions that show task-related increases in activity during externally directed attention (Buckner et al., 2019. Shulman et al., 1997). These task positive regions have been referred to as the task positive network (TPN), which typically includes dorsal and frontoparietal cortical regions, or the dorsal attention network (DAN) and frontoparietal network (FPN) (Spadone et al., 2015. Buckner et al., 2019).

This anti-correlated relationship between the DMN and TPN has been implicated in attentional control, and DMN FC is disrupted in pathologies such as attention deficit hyperactivity disorder (Uddin et al., 2008. Konrad et al., 2010. Liddle et al., 2011. Hale et al., 2014. Hoekzema et al., 2014. Jansen et al., 2017. Abbas et al., 2019b, Bauer et al., 2020). Increased default mode network activity has also been associated with resting state mind-wandering (Mason et al., 2007. Andrews-Hanna et al., 2010. Golchert et al., 2017. Mittner et al., 2016) and increased rumination, both thought to be predictors of unhappiness (Smallwood et al., 2009. Killingsworth et al., 2010).

However, while increased DMN activity may drive spontaneous thoughts in the form of mind-wandering and maladaptive rumination, it may also signal an increase in all types of spontaneous cognition, including creativity (Christoff et al., 2016). Several groups have reported a link between increased activity in the primary nodes of the DMN and creativity, including mPFC and PCC (Fink et al., 2010. Liu et al., 2012. Beaty et al., 2015). An increase in functional connectivity between the DMN and the frontoparietal/executive control network (FPN) and the DMN and ventral-attention/salience network (VAN) has also been implicated in creativity, specifically in the domain of musical improvisation (Bengtsson et al., 2007. Limb et al., 2008. Loui, 2018. Beaty et al., 2015). Such alterations in DMN-FPN or DMN-VAN connectivity may represent an increased capacity for flexible cognitive control driving creativity (Zabelina et al. 2018. Li et al., 2021).

Based on the known relationship between DMN-FPN FC and creativity, Belden et al., 2020 explored changes in resting state functional connectivity (FC) depending on creative musical training, using seed-based functional connectivity and graph-theory analysis to compare changes in the DMN, FPN, and other networks. Belden et al., 2020 found that improvisational musicians showed an increase in resting state FC between the visual network and both the DMN and FPN, as well as increased intrinsic FC in the ventral DMN when compared to classical musicians and controls.

While traditional time-averaged functional connectivity analysis employed by Belden et al., 2020 can identify seed-based correlations during resting state fMRI, analysis of dynamically recurring patterns such as co-activation patterns or infraslow quasi-periodic patterns (Majeed et al., 2011. Petridou et al., 2013. Liu et al., 2018. Yousefi et al., 2018. Yousefi et al, 2021. Bolt et al, 2022. Meyer-Baese, Watters, 2022) may provide deeper insight into the relationships between networks involved in creative improvisation. In this study, we seek to expand upon the network based FC and graph theory analysis made by Belden et al., 2020 by applying dynamic functional connectivity methods to the same dataset, consisting of classical musicians (classical), improvisational musicians (improv) and minimally trained/non-musicians (MMT).

Multiple types of recurring dynamic patterns have been detected and described in rsfMRI data (Hutchison et al., 2013. Keilholz et al., 2017. Preti et al., 2017. Lurie et al., 2020. Meyer-Baese, Watters, 2022). The present study focuses primarily on the dominant resting state pattern of anti-correlated activity between the DMN and the task positive regions, which occurs quasi-periodically on the infraslow timescale (with 1 cycle lasting about 20 seconds in humans) (Abbas et al., 2019a). This pattern, which has been described as the primary quasi-periodic pattern (QPP) of dynamic FC, was first quantitatively detected in blood-oxygen level dependent (BOLD) fMRI in anesthetized rats (Majeed et al., 2009), and was subsequently detected in human rsfMRI (Majeed et al., 2011). The original QPP detection in Majeed et al., 2009 and Majeed et al., 2011 employed a sliding window based algorithm that converged on a reliable pattern of DMN/TPN anti-correlation (Majeed et al., 2009. Majeed et al.,2011). More recently, this QPP has been replicated using complex principal components analysis (cPCA) on rsfMRI (Bolt et al., 2022), and appears to be one of the three dominant spatiotemporal patterns that account for most of the time lagged connectivity structure in intrinsic brain activity observed across methods.

Given the relation of default mode QPP dynamics to attention, we predicted that prolonged focused musical training, like that experienced by classical musicians, may lead to altered DMN and DAN anti-correlation dynamics when compared to musicians who are primarily improvisational. The established role of the DMN in spontaneous cognition makes it a useful reference network for making functional correlations that may implicate other networks in creative cognition. We primarily measured the amount of correlation between other canonical networks and the DMN at the group and individual level, over the full cycle of the QPP. QPP analysis allows us to capture whole brain dynamics and the time lag structure between other cortical networks based on musical training, and to compare with the inter-network results originally reported in Belden et al., 2020 in networks such as the visual network and frontoparietal network. Thus, while our analysis was primarily focused on exploring functional correlation to the DMN during the QPP, we also followed QPP results that differed in regions beyond the DMN.

In addition to QPP detection using the QPP-finding algorithm, we repeated part of our group level QPP network analysis using cPCA, a type of dimensionality reduction method originally pioneered in geological and climate sciences to detect the components that explain most of the variance within propagating patterns (Horel, 1984). More recently, Bolt et al., 2022 demonstrated that in global signal regressed data, the first component from cPCA in rsfMRI is equivalent to the QPP of DMN/TPN activity. To our knowledge, this is the first study to apply both the pattern-finding algorithm and cPCA to the same dataset to detect group-wise dynamic FC differences. By applying both methods, we hoped to uncover any differences in the sensitivity of each approach to group level changes in QPP network dynamics. To compare group-averaged vs individual QPP network trends, we repeated all algorithm-based QPP detection on a subject-wise level.

Using these two methods, a QPP-finding algorithm and cPCA-based QPP detection, we hypothesized that: (1) DMN network activity in the QPP may be higher in improv musicians than improv or MMT subjects, given the association between DMN activity and mind-wandering/spontaneous cognition, (2) the visual network in the QPP would be more strongly correlated with the DMN and FPN, given the seed-based connectivity results in Belden et al., 2020, (3) group level differences in QPP network correlations would be consistent across both the sliding window and cPCA methods, (4) the application of QPP waveform analysis may detect groupwise differences in time-lagged information missed by traditional seed-based FC analysis.

The findings in this study deliver a better understanding of how creativity, a type of spontaneous cognition, alters the default mode network’s dynamic interaction with other major cortical and sensory networks, ultimately furthering our understanding of the complexities of musical training, creativity, and brain network dynamics.

## 2. Methods

All code is Open-Sourced and available online. QPP code and cPCA code referenced in the methods section is available on github: (1) https://github.com/BnzYsf/QPP_Scripts_v0620, and (2) https://github.com/tsb46/complex_pca

All code used for sliding window quasi-periodic pattern detection was run in Matlab (Mathworks Inc. 2023). Visualizations were generated through R (RStudio Team, 2023) and FSL (Jenkinson et al., 2012). Code for cPCA was run in Python on a Linux operating system.

### 2.1 Participants

Structural and functional MRI scans were obtained from Alex Belden and Psyche Loui at Northeastern University. All original scan acquisition adhered to best practices for ethical human data collection and informed consent in accordance with local institutional review boards. For a full description of participants and MRI acquisition see their original study Belden et al., 2020.

In brief, 48 young adult subjects were recruited from universities and schools in the Boston area and classified into three groups based on musical training background: classical training, improvisational training, or minimal-musical training (MMT). Final groups consisted of 16 subjects each (n = 4 females, n = 12 males). To the extent possible, the groups were matched in age, general cognitive ability, pitch discrimination, duration of musical training, and age of onset of musical training.

### 2.2 Data acquisition and pre-processing

T1 weighted structural scans and resting state functional scans were obtained on 3T Siemens scanners at Northeastern Biomedical Imaging Center and the Olin Neuropsychiatry Research Center. T1-weighted sequences were 3D magnetization prepared rapid-acquisition gradient-echo (MPRAGE) with a voxel size of 0.8 x 0.8 x 0.8 mm^3^ (TR = 2.4 s, TE = 2.09 ms, flip angle = 8°, FOV = 256 mm).

Resting state scans had a duration of 7.5 minutes and were obtained with an echo-planar imaging (EPI) sequence with 947 volumes (TR = 475 ms; TE = 30 ms; flip angle = 90, 48 slices; FOV = 240 mm; acquisition voxel size = 3 × 3 × 3 mm^3^). Following typical resting state protocol, participants were instructed to keep their eyes open and fixated on a cross for the duration of the scan.

For processing, scans were formatted according to Brain Imaging Data Structure (BIDS: https://bids.neuroimaging.io/. Poldrack et al., 2024). All pre-processing was run in Linux (Ubuntu 22.04.3 LTS) using the configurable pipeline for the analysis of connectomes (C-PAC: https://fcp-indi.github.io/). Outputs for algorithm based QPPs were generated with and without global signal regression (GSR); non-GSR results are available in supplemental material. As part of the default C-PAC pipeline, anatomical scans were registered to the 2mm Montreal Neurological Institute MNI 152 Atlas. Resting state functional scans were also registered to standard MNI space and then extracted as timeseries to the Brainnetome 246 atlas, a 246 region-of-interest (ROI) parcellation based on MNI space. The first 10 volumes of each scan were truncated for all subjects, resulting in a final 937 timepoints by 246 ROI 2-dimensional timeseries for each subject.

### 2.3 Quasi-Periodic Pattern Acquisition window

Recurring spatiotemporal patterns (quasi-periodic patterns) were identified using an updated version of the sliding window pattern detection algorithm from Majeed et al. 2011 and other work from the Keilholz group. For a more detailed description of the original algorithm see Majeed et al. 2011. Previous versions of the algorithm were based on a user-defined or random starting point within the time-series, and would conduct a sliding correlation of the initial segment with other segments in the scan that exceeded a threshold of correlation (0.2) with the initial template. In this study, we used a robust version of the pattern algorithm that starts at the beginning timepoint and then iterates through each timepoint, updating until a convergent pattern is produced (Yousefi et al., 2018 Xu et al., 2023).

While the QPP algorithm we used no longer has a user defined start point, it still requires a user defined window length. Previous work indicates that QPPs last approximately 20s in humans (Abbas et al., 2019a. Bolt et al., 2022), but we ran the QPP algorithm with various window lengths (WL) to determine empirically which WL most reliably captured a full phase of the QPP in this dataset. After trying a range of window lengths between 15-40 seconds, we found that a WL of 24 time points most reliably captured a full phase of the QPP in this dataset and used that WL for all algorithm based QPP analysis. Given that the TR of the functional scans was .475 seconds, this means that the QPPs displayed in the results section are on the order of 12 seconds (TR x WL, .475 seconds X 24 second WL = 12 seconds).

Initial QPP analysis was conducted at the group level: we ran the QPP algorithm on the concatenated timeseries for 16 subjects at a time (classical, improv, MMT, respectively) and an average group level QPP was detected. QPP analysis was then conducted between group-level QPPs based on canonical networks (described below). Individual QPPs were also run for all 48 subjects. Statistical comparisons at the individual level were only made for the network correlations found to be most different at the group level, including the visual network and parts of the subcortical network (amygdala).

### 2.4 Network based analysis of QPP functional connectivity

Analyzing QPP dynamics with canonical network definitions generated results that could be easily compared with the default mode network literature as it relates to creativity, including Belden et al.’s original seed-based analysis findings (Belden et al., 2020). We therefore decided to employ canonical network definitions after applying the QPP pattern detection algorithm.

After QPP detection, the QPP was mapped to network space (Yeo et al., 2011). ROIs from the Brainnetome 246 parcellation were assigned into 7 canonical cortical networks plus subcortical, resulting in QPP waveforms for the following 8 canonical networks: default mode network (DMN), frontoparietal network (FPN), dorsal attention network (DAN), ventral attention network (VAN), somatomotor network (SOM), visual network (VIS), limbic network (LIM), and subcortical network (SCN).

From these timeseries, waveform plots were then generated plotting the normalized BOLD signal for one full cycle of the QPP for all defined networks. We then compared the correlations, the amplitude, phase, and squared difference between networks of each group-level QPP against all other groups. As the present study was focused on dynamics relative to the DMN, the focus of these comparisons was with respect to the DMN. As the improv group showed an altered amygdala-DAN FC, we reported differences with respect to the DAN instead of the DMN. Because the DMN and DAN are strongly anti-correlated in all group QPPs either the DMN or DAN provide convenient reference networks to compare the remaining Yeo’s cortical networks.

For statistical measures, group level network correlation comparisons were made using the cocor R library (Diedenhofen et al., 2015). Subject-wise statistical comparisons of network correlations were made in R studio using Kruskal-Wallis test for multiple comparisons from base R studio statistical packages (RStudio Team, 2023). All correlation values were Fisher transformed. Subject-wise statistical comparisons were made after Fisher transform. Correction for multiple comparisons was made using the conservative Bonferroni correction (VanderWeele and Mathur, 2019) based on the assumption of 7 underlying comparisons between the 8 canonical networks used. Thus, the standard significance threshold of α = .05 was adjusted for 7 comparisons (.05/7) to an α = .007.

### 2.5 Complex Principal Components Analysis

Bolt et al, 2022 demonstrated that the majority of variance in low-frequency spatiotemporal BOLD patterns can be explained by three principal components (Bolt et al., 2022). In global signal regressed data, the first of those principal components is equivalent to the QPP or task-positive vs default mode anti-correlated pattern. Note that in Bolt et al. the primary results are reported without global signal regression, and thus in that study the first component is global signal while the second component is equivalent to the QPP. Using the same methodology of Bolt et al., we applied Complex Principal Components Analysis (cPCA), a dimensionality reduction method, to identify the principal component of low-frequency BOLD signal in our dataset equivalent to the QPP.

The same 2-dimensional timepoint by ROI preprocessed CPAC outputs that were the QPP algorithm input were used for cPCA. cPCA was run in Python on a Linux operating system on concatenated scans on a groupwise basis for the 3 respective music groups (classical, improv, MMT).

cPCA was run initially with 10 components; consistent with Bolt et al., 2022, the majority of variance in our dataset was explained by the top 3 components (Figure 4A). We then plotted the top 3 components across 937 timepoints to generate histograms of the proportion of the components across subjects (Figure 4B) and total proportion of the top 3 components (Figure 4C). Note that components 2 and 3 of GSR data possibly correspond to additional types of QPPs, QPP2 and 3 specifically (Yousefi, 2021), but as the present study focuses on the dominant QPP of DMN/TPN anti-correlation the other QPPs/components were not used for analysis.

After confirming that the relative proportion of time spent in the QPP component was similar across groups (Figure 4B-C), component 1 (QPP1) for each musical group was then reconstructed as a 4-dimensional nifti file for visualization in FSLeyes. We then plotted the QPP component as a reconstructed time series into brain space and generated network activity waveforms for comparison to the sliding window algorithm results (Figure 5).

cPCA based waveforms were generated in FSLeyes (Jenkinson et al., 2012) by selecting voxels in key nodes of ROIs corresponding to each network of interest (DMN, DAN, VIS, AMYG) based on the Brainnetome parcellation (table 1) and then plotting the timeseries in bins representing the length of one full cycle through the QPP. The default number of bins is 30 for the reconstructed time series. 100 bins were used instead for increased temporal resolution for plotting the phase aligned components. For any number of bins used in the reconstructed component time series, the total number of bins represents one full cycle through the QPP, corresponding to radial distance between 0 and 2*π*. Similar to the algorithm based QPP, one full cycle of the cPCA based QPP takes approximately 20 seconds in humans (Bolt et al., 2022). Note that in the algorithm based QPP we used a window length of 24 time points, which was determined empirically by trying a range of window lengths between 15-40 time points.

**Table 1:** MNI and Brainnetome coordinates for all seeds used to generate cPCA waveforms.

| Network | MNI coord. left hemisphere | MNI coord. right hemisphere | Brainnetome ROI | Brainnetome ROI numbers |
| --- | --- | --- | --- | --- |
| DMN | 87 85 107 | 95 85 107 | PCC | 175,176 |
| FPN | 70 151 124 | 113 151 124 | Dorsolateral area 8 | 3, 4 |
| DAN | 72 125 139 | 110 125 139 | Dorsolateral area 6 | 7, 8 |
| VIS | 77 26 63 | 107 26 63 | Occipital polar cortex | 203, 204 |
| AMYG | 70 125 51 | 114 125 51 | Medial amygdala | 211, 212 |

As with the algorithm based QPP networks, we then made group-level comparisons of the correlations between the networks of interest and the default mode network. For the cPCA QPP analysis, only group level comparisons were made, no subject-wise waveforms or correlation values were generated. When plotting ROI based correlation of cPCA based time courses in FSLeyes, we found that all voxels within a given Brainnetome ROI were highly correlated, near 1 (see supplemental figure 2), and thus plotting the time course from a representative voxel within each ROI was roughly the same as plotting a mask for each ROI. Thus, for each network of interest a representative voxel was selected and used to generate waveform plots (Figure 5 B-C). To account for interhemispheric differences, voxel-based time courses were plotted from both left and right hemispheres. No qualitative differences were noted in left vs right hemisphere time courses from the chosen voxels, so the average time course between left/right was calculated and plotted to compare with QPP pattern algorithm results (Figure 5 B). All voxels and their corresponding Brainnetome ROI numbers are shown in table 1.

## 3. Results

### 3.1 Network analysis of QPPs

QPPs detected in our groups were similar to those in prior studies (Abbas et al., 2019a. Bolt et al., 2022), showing a strong pattern of anti-correlation between default mode and dorsal attention/task-positive areas (Figure 2). Group level QPPs are discussed first followed by a comparison with subject-wise QPP results.

**Figure 1:**
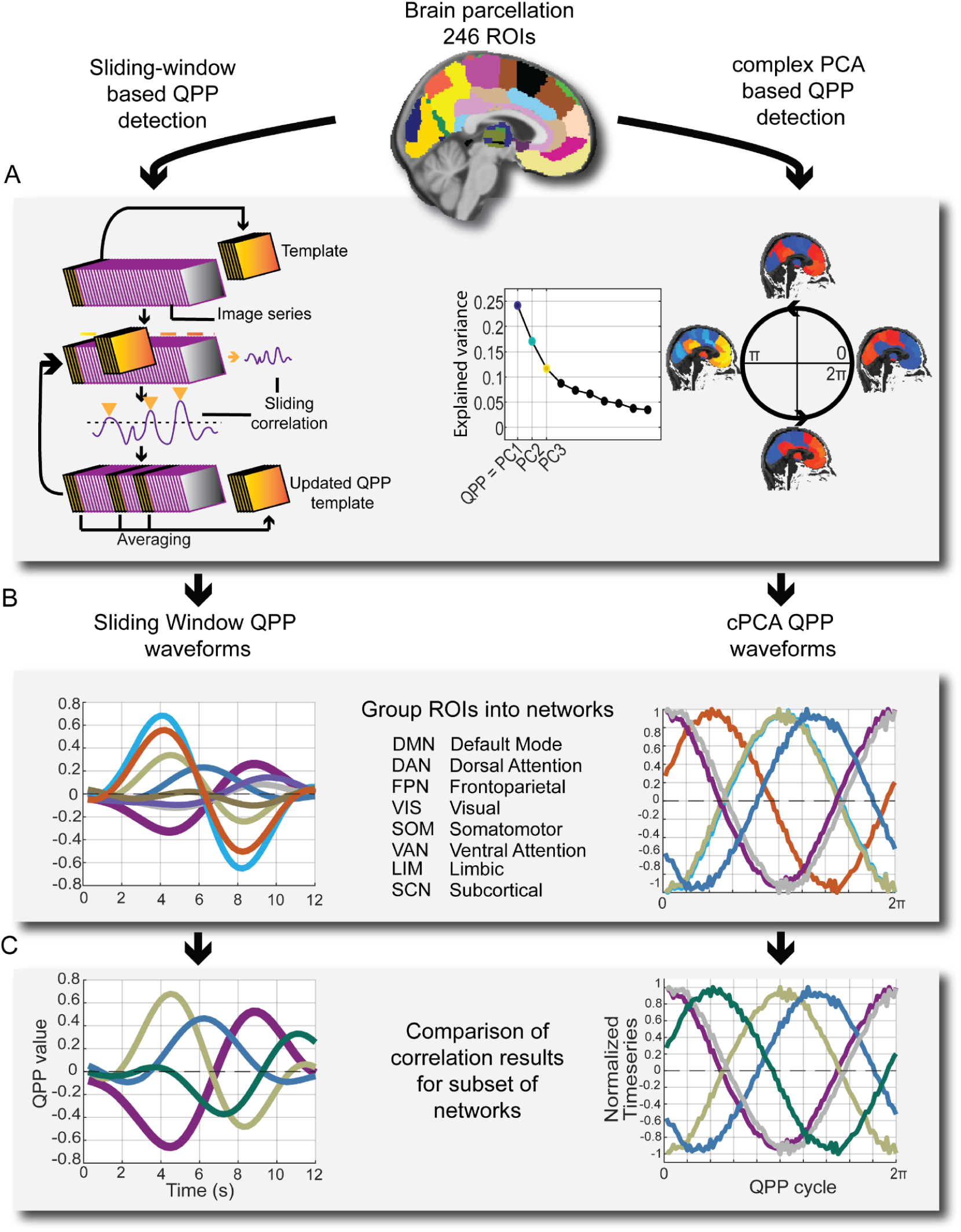
Parcellation of pre-processed resting state fMRI time series into Brainnetome 246 ROIs. A) Detection of quasi-periodic patterns (QPPs) using a sliding window based algorithm and complex principal components analysis, respectively. B) Grouping of 246 ROIs into 8 canonical networks for C) comparing correlations and squared differences of network activity during the QPP.

**Figure 2:**
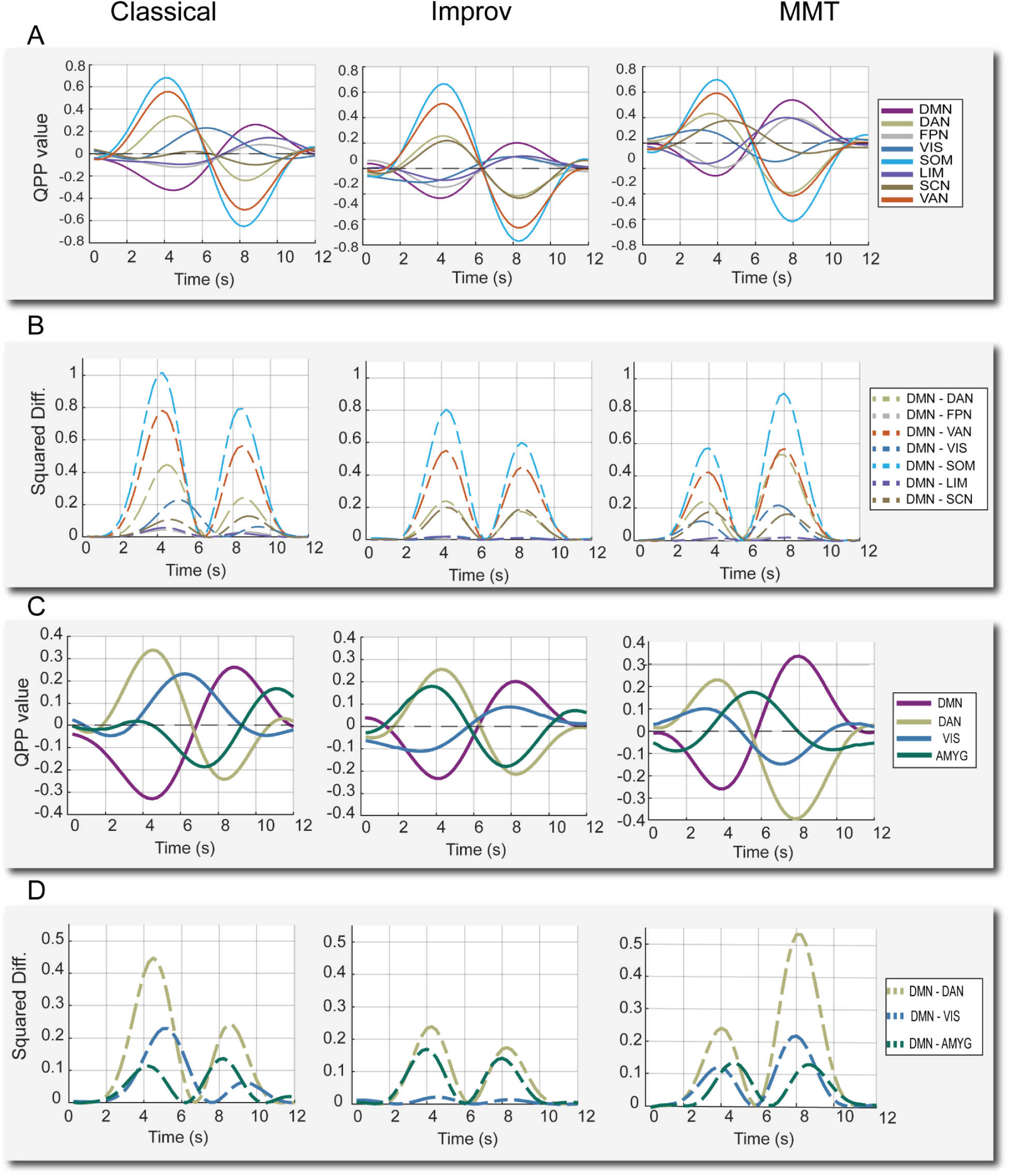
QPPs from sliding window pattern detection algorithm. A) QPP waveforms during 1 full cycle for all 8 cortical networks. DMN: default mode network, DAN: Dorsal attention network, FPN: frontoparietal network, VIS: visual network, SOM: somatomotor network, LIM: limbic network, SCN: subcortical network, VAN: ventral attention network. B) Squared difference between each network and the DMN for the same cycle. C) QPP waveforms for the DMN, DAN, and the two regions where group-level differences were noted in improv musicians: the visual network and amygdala. D) Squared difference between the same subset of networks for each group.

DMN/TPN anti-correlation is shown to be associated with changes in cognition and arousal, specifically attentional control and mind wandering, (De Havas et al., 2012. Belloy et al., 2018. Abbas et al., 2019, Godwin et al., 2017. Golchert et al., 2017. Chou et al., 2022. ). Given the prolonged hours of repetitive musical training undergone by classical musicians, we initially speculated there may be differences in resting attentional control and thus looked for differences in correlation between default mode and task positive regions (DAN, FPN, and VAN) in classical musicians. However, we did not find significant differences in correlations specifically between the DMN/TPN regions in the classical musicians (Figure 2 A). Instead, we found significant differences in the time lagged network correlations between the DMN and other networks in the improv group. Additionally, the amplitude of QPPs in improv musicians trended much lower than classical and MMT groups.

As captured by the squared difference between networks, the overall amplitude of the anti-correlation between the DMN-DAN in the improv group trended far below the classical and minimally trained musicians (Figure 2, C-D). For the improv musicians group, the squared difference between DMN-DAN was 0.076, roughly half of the classical group’s 0.132 and the MMT group’s 0.149 (note that squared differences between normalized network activities were measured at the group level and thus lack a standard deviation).

In addition to changes in overall QPP amplitude, the improv group also showed an altered QPP progression in the visual and subcortical networks. More specifically, the network waveform plots (Figure 2C) and their underlying correlations showed increased correlation between visual-DMN and decreased correlation between amygdala-DMN activity in the improv group QPP (Figure 3B. Given the strong anti-correlation between the DMN and DAN in the QPP, decreased amygdala-DMN FC also means increased amygdala-DAN FC. As previous literature implicates amygdala-DAN FC as an area of interest for behavior and cognitive control (He et al., 2015, Sylvester et al., 2020), we decided to report the difference in extrinsic amygdala FC as between amygdala-DAN and not amygdala DMN. We found no clear group-wise differences in other subcortical structures such as the hippocampus, which was highly varied between groups, or in the thalamus and basal ganglia, which seem to show almost no QPP-like activity in this frequency range (0.1-1 Hz). This is consistent with Yousefi et al., 2021, which reported very weak thalamic QPP activity only revealed by substantial averaging (Yousefi et al., 2021).

**Figure 3:**
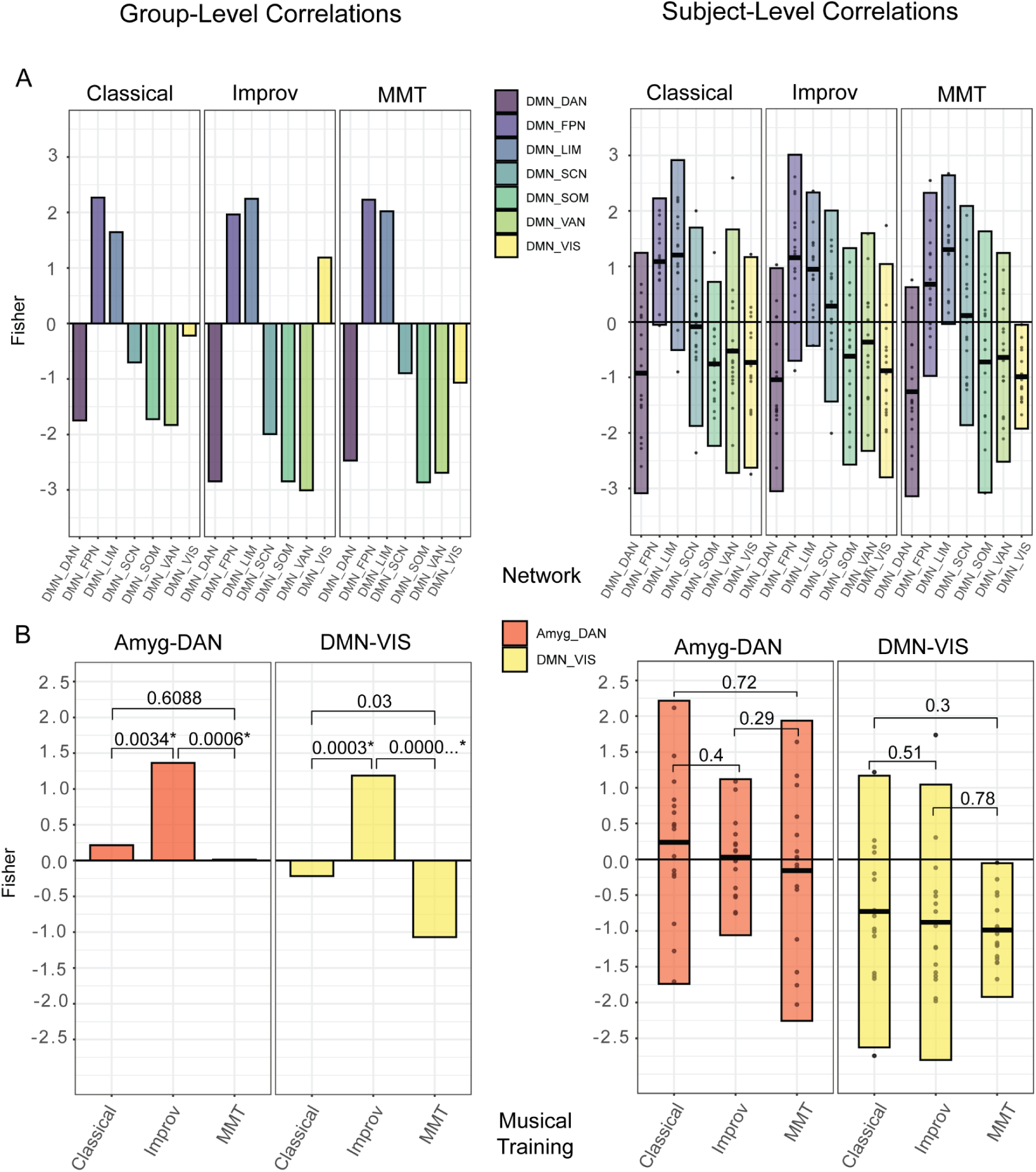
A) group (left) and individual (right) QPP correlations between all networks and the DMN. B) Group and individual QPP correlations only between the Amyg-DAN and DMN-VIS, respectively. The improv group showed significantly higher FC between Amyg-DAN and DMN-VIS compared to classical and MMT. Group level p-values generated with cocor tool (Diedenhofen and Musch, 2015), individual level p-values generated with Kruskal-Wallis test for multiple comparisons in R studio. A Bonferroni adjusted significance threshold of .007 was used for both subject-wise and group comparisons, assuming a multiple comparison correction of .05/7 (assuming 7 network comparisons between the 8 networks defined).

Given that the improv group’s visual network and amygdala seemed to be the most different, plots and statistical comparisons were made for the improv group’s visual network-DMN and amygdala-DAN correlations compared to the other training groups. The improv musicians showed an increased correlation between both the visual network and DMN and the amygdala and DAN relative to the classical and minimally trained musicians (Figure 3, A-B, left).

Group-level QPP waveforms were also generated on time series processed without global signal regression (GSR). Waveform results without GSR exhibited positive global correlation between all of Yeo’s networks used in the present study (supplemental Figure 1).

### 3.2 Subject level QPPs for comparison of selected networks

Group level analysis is commonplace in seed-based and dynamic FC studies but obscures individual variability. To explore how dynamic DMN FC differs at the subject-wise level, we also applied the QPP algorithm individually to all 48 subjects. We then again plotted the fisher-transformed correlation values between the DMN and all other networks for each subject (Figure 3).

The purpose of the individual QPP comparison was two-fold. First, we wanted to know how similar the mean network correlation values of all networks in the individual QPPs were to the correlations detected at the group level. Previous work from Yousefi et al., 2018 using neurotypical subjects from human connectome data suggests that DMN/TPN anti-correlation is fairly consistent within individual subjects, and between the individual and group level. However, Yousefi et al., 2018 did not apply QPP analysis to detect group-wise differences based on training or pathology, and focused on DMN/TPN without considering other networks. Having individual correlation values allowed us to more easily make statistical comparisons as they provide a distribution and variance.

We found notable differences between the group and subject level QPP analysis. First, at the group level (Figure 3, A-B) the improv musicians showed a much higher correlation between the DMN and visual network (Fishers r = 1.187) than classical (Fishers r = -0.218) and non-musicians (Fishers r = -1.068993262). This DMN-VIS correlation increase is not present in the subject level analysis. Second, improv musicians also showed a significantly higher Amyg-DAN correlation at the group level (Fishers r = 1.365) compared to classical (Fishers r = 0.215) and non-musicians (Fishers r = 0.0142). Once again, when the QPP algorithm was run on a subject-wise basis this increased Amyg-DAN correlation was not present (Figure 3 B).

### 3.3 Complex Principal Components Analysis of QPPs

As in Bolt et al., 2022, the top 3 components explained the majority of the variance and there is a strong elbow in the explained variance after the first 3 components (Figure 4A). We found the proportion of the top 3 components across all 937 timepoints for each subject and group to be very similar (Figure 4 B-C). For all 3 training groups, the first component (representing the QPP) was the dominant brain state for approximately 40-50% of all timepoints (Figure 4C). Note that the other components (2-3) were not used for further comparison as they represent different spatiotemporal patterns (Bolt et al., 2022).

**Figure 4:**
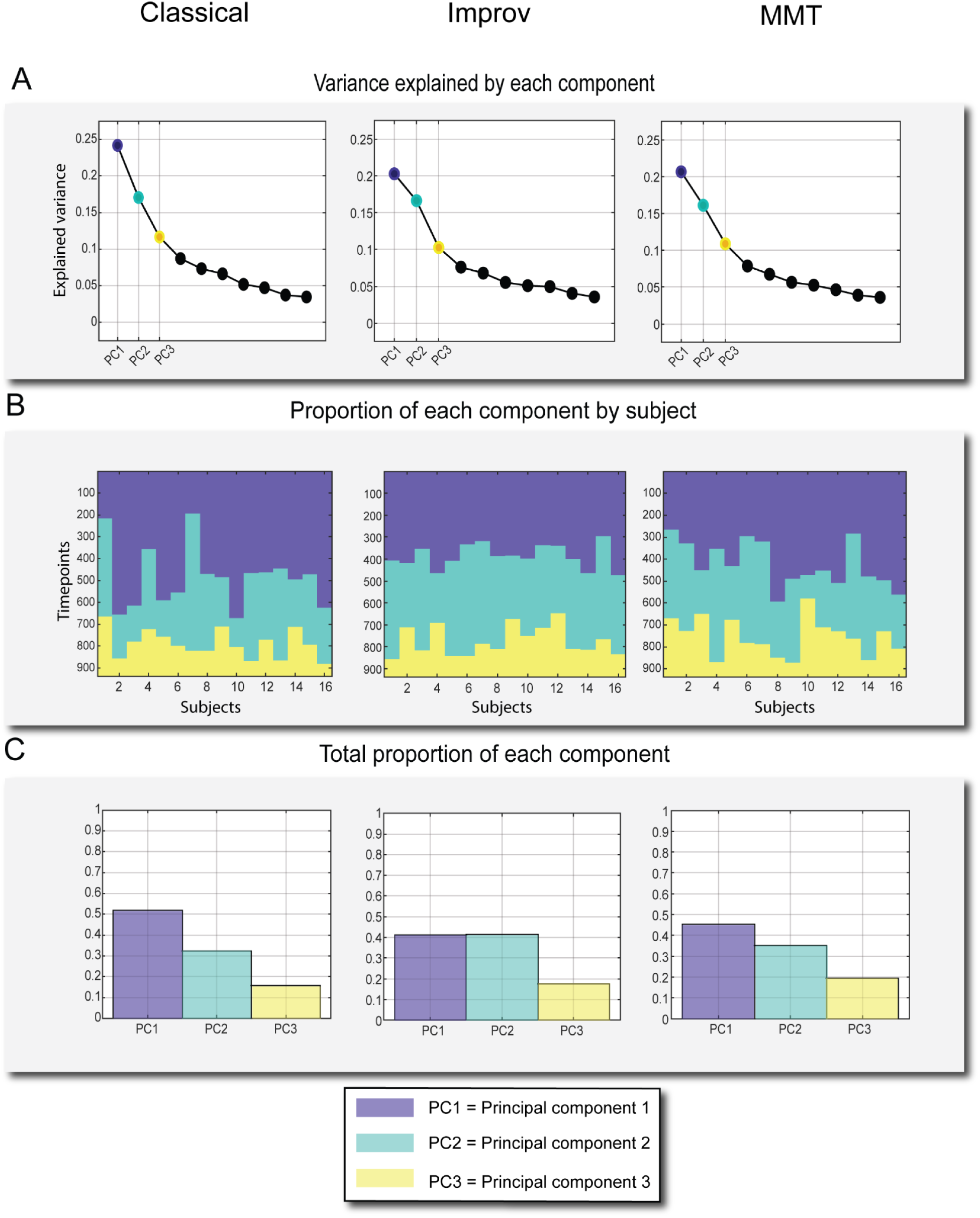
Explained variance and occurrence of top three components from complex PCA. A) Explained Variance of the top 10 components, top 3 highlighted. B) Top 3 components’ respective proportions plotted against 937 timepoints. C) Proportion of each of the three components over the course of the scan out of 1.

The cPCA results are aligned in several ways with the QPP algorithm results. First, the peak amplitude of the QPP component is noticeably less in the improv group, roughly half that of the other training groups, just as it was with the DMN-DAN amplitude in the algorithm based pattern detection. Specifically, improv musicians’ normalized cPCA timeseries (BOLD signal) showed a decreased peak amplitude range in the QPP of -0.462 to 0.462, compared to the classical musicians’ range of -0.816 to 0.816 and the MMT range of -0.772 to 0.772 (Figure 5A). Second, when mapping the cPCA-based QPP back to brain space we noticed that in the improv group, the occipital/visual regions were positively correlated with the posterior cingulate cortex (r = 0.371), a primary node of the default mode network (Figure 5A). The classical (r = -0.549) and MMT (r = -0.960) groups showed typical visual-DMN anti-correlation, consistent with the sliding window QPP waveforms. Thus, based on the reconstructed time series in brain space, the improv musicians showed increased visual-DMN correlation just as they did in the QPP algorithm approach. Additionally, the improv musicians also showed a much higher correlation between the amygdala and DAN (r = 0.878) compared to the classical (r = 0.215) and MMT (r = 0.014) groups, again consistent with the QPP algorithm results (Figure 5 B-C).

**Figure 5:**
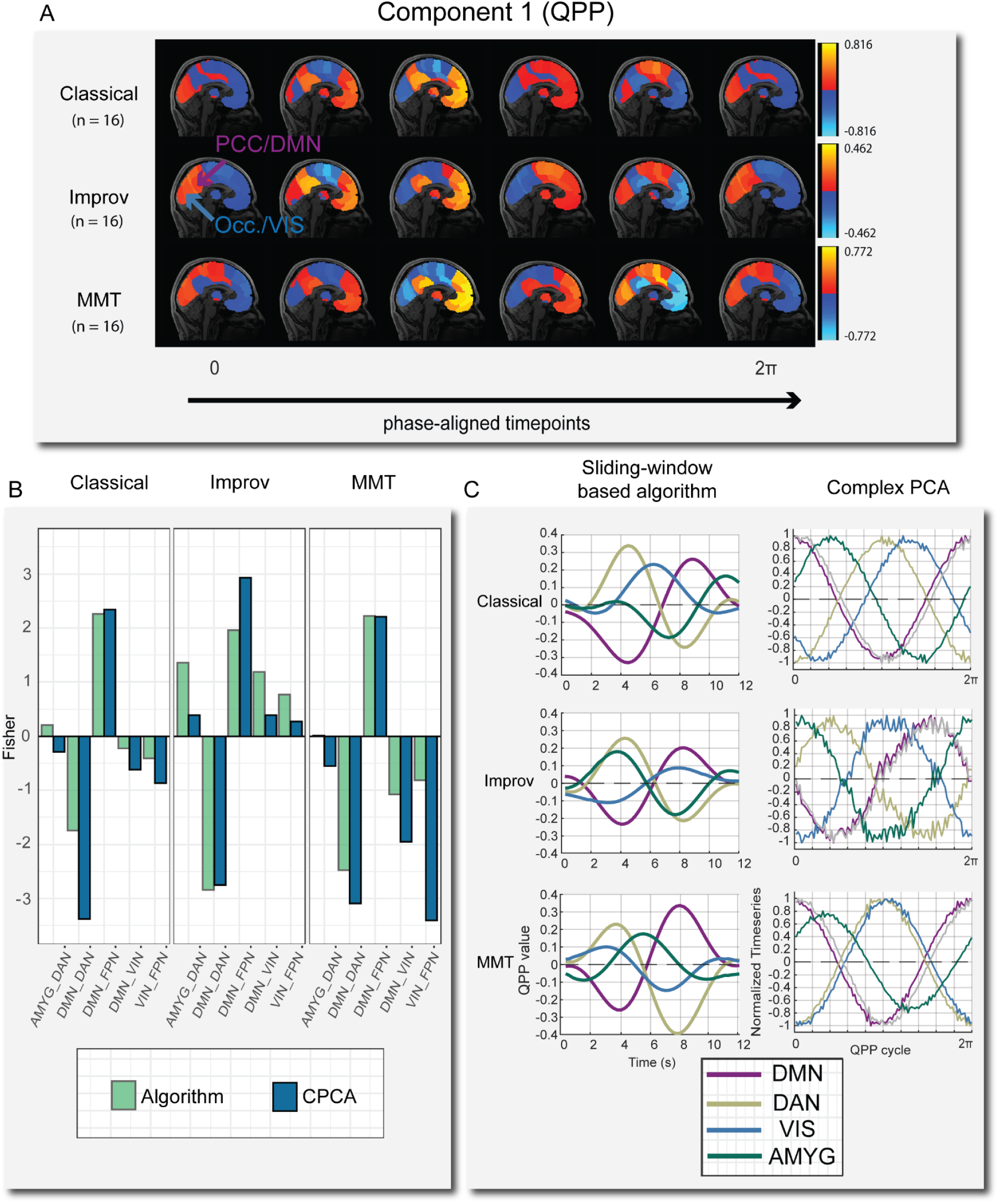
A) One full phase (zero to 2π) of group-level cPCA detected QPP time-courses mapped onto brain space. Time-courses were manually phase adjusted into 6 time points. Note that in the improv group the posterior-cingulate cortex (PCC) is highly correlated with primary visual cortex (occipital lobe) at various time points, where in the classical and minimally trained musicians these two regions are always anti-correlated. B) Comparison of group-level correlation results between the sliding window-based algorithm and cPCA for selected networks. The overall network trends are very similar in both approaches. C) Waveform plots of normalized BOLD activity from QPP algorithm (left) and cPCA approach (right) for one full cycle of the QPP.

In summary, we found that the three main group level differences detected in improv musicians were consistent between the pattern detection algorithm and complex PCA methods. (1) The overall amplitude of the QPP between DMN-DAN trends much lower in the improv group. (2) The visual network in the improv musicians has a positive correlation with the default mode network at the group level, whereas for the classical and MMT groups the visual network-DMN were anti-correlated. 3. The amygdala is more correlated with the dorsal attention network in the improv musicians than in classical musicians or in MMTs.

## 4. Discussion

This study applied two methods of dynamic functional connectivity analysis to a dataset of human subjects based on musical training. We found convergent results at the group level from the sliding window based QPP algorithm and complex PCA approaches. In particular, both dynamic FC analysis methods detected an increased FC between the visual network and both the DMN and FPN, respectively, in improv musicians compared to classical and untrained musicians. This increased VIS-DMN and VIS-FPN connectivity is consistent with the previous group level findings from Belden et al., 2020, where network based FC was conducted using traditional seed based FC analysis and graph theory. Notably, the DMN-VIS results were also robust across different atlases, given that Belden et al., 2020 did not use the Brainnetome atlas as we did. Unexpectedly, we also detected increased FC in the improv musicians between the medial amygdala and DAN with both dynamic FC methods. With both methods, a trend of decreased QPP amplitude was detected in the improv group. When repeating QPP detection with the sliding window algorithm at the subject-wise level, the group-level differences were not replicated. The possible implications for these findings are discussed below.

### 4.1 Default Mode - Visual Network Connectivity and Creativity

In both QPP detection methods the visual network/occipital lobe activity of improv musicians was more in phase with the DMN and FPN compared to classical and MMT groups, resulting in a significantly higher VIS-DMN and VIS-FPN correlation. This finding is consistent with the seed based connectivity analysis employed by Belden et al., 2020. The robustness of this finding across two dynamic QPP methods, in addition to the original static FC analysis, seems to indicate that an increased VIS-DMN/FPN connectivity is a very strong feature of this dataset. Whether increased VIS-DMN connectivity is a real feature of improvisational musical training or related to an increased capacity for creativity is a much more difficult question to answer conclusively with these findings alone.

However, these dynamic FC results contribute to a growing body of evidence indicating that visual-DMN connectivity may drive domain general creativity by way of increased mental imagery (Belden et al., 2020). A link between visual network activity and creativity was originally supported by findings from EEG (Petsche, 1996) and white matter tractography (Takeuchi et al, 2010. Zamm et al., 2013). For example, Petsche found increased coherence between frontopolar and occipital/visual regions during visual, verbal, and musical acts of creative thinking (Petsche, 1996). More recent rsfMRI findings also indicate that spontaneous visual network activity and connectivity to the DMN are related to visual creative cognition (Chen et al., 2019. Wang et al, 2021).

Considering the role of activity in V1 and V2 in the generation of mental imagery (Pearson, 2019), it seems intuitive that creativity may be linked to visual imagination. However, the combination of visual imagination and musical creativity, a process presumably dominated by audition more than vision, is less intuitive. At the same time, visual activity has already been linked to spontaneous imagery and creativity in non-waking states of consciousness. Dreaming during REM sleep for example, another type of spontaneous cognition, is heavily linked to both default mode network (Christoff et al., 2016) and spontaneous visual network activity (Eagleman and Vaughn, 2021). Dreaming itself has also been linked to creativity (Schredl 2007. Barrett, 2017). Given the available evidence, it seems reasonable that increased synchrony between two networks heavily implicated in the generation of spontaneous thoughts could explain an increased capacity for domain general creativity.

Of course, even if the increase in dynamic VIS-DMN FC in this dataset is related to creative cognition, and not merely an artifact, the direction of causality is unclear. In other words, from the analysis in this study it is unclear whether years of creative musical improvisation cause an increase in VIS-DMN connectivity or whether people who are more creative and exhibit such dynamic FC in the first place are simply more drawn to certain types of creative expression.

### 4.2 Dorsal Attention - Amygdala Connectivity

The improv group QPP also displayed increased correlation between the amygdala and DAN for both QPP detection methods. It is difficult to make strong interpretations of increased amygdala-DAN connectivity based on the analysis from this study alone. However, there is related evidence from rsfMRI studies that an increased amygdala-DAN correlation could underlie important behavioral and attentional changes. For example, one rsfMRI study using the same canonical networks from the present study (Yeo’s 7 networks) found that increased amygdala-DAN FC was negatively correlated with trait anxiety (He et al., 2015). If our results of increased amygdala-DAN functional connectivity were to be replicated in other datasets measuring creativity, this would establish a potential link between creativity training and anxiety.

More recently, Sylvester et al., 2020 empirically defined three amygdala subsections based on their respective functional connectivity with major cortical networks: identifying a default mode subdivision, dorsal attention subdivision, and a subdivision that did not display a preferred connectivity with the DMN or DAN. In the present study, the amygdala section used would have fallen within what Sylvester et al., 2020 described as the centromedial amygdala, or DMN connected amygdala subdivision. However, in our improv musicians, despite the selected amygdala seed being in the DMN-associated region, the amygdala was instead highly correlated with the DAN. In contrast, the classical and MMT groups displayed positive DMN correlation from the same amygdala voxel (or DAN anti-correlation), consistent with the neurotypical trend described in Sylvester et al., 2020. Again, the direct implications for this altered amygdala functional connectivity are unclear. But as Sylvester et al., 2020 points out, the subsections of the amygdala with high DAN functional connectivity may regulate top-down attentional and spatial processing (like that typically associated with the DAN). If increased amygdala-DAN FC is a real feature of increased creativity or improvisation, it could be the case that the amygdala’s role in modulating the DAN becomes altered in creatively trained subjects.

There is also some anatomical evidence that the amygdala is related to increased creative cognition, with Bashwiner et al., 2016 reporting that increased volume in the left amygdala was significantly correlated with higher scores of creativity in their subjects. In a study specifically using jazz musicians and improvisation, the amygdala showed increased activity during musical improvisation inspired by positive images (McPherson et al., 2016). Whether increased amygdala activity or amygdala-DAN FC at rest could signal increased creativity will require further replication in future studies.

### 4.3 Group-wise vs subject-wise results

Differences in functional connectivity results between group-averaged and individual level analyses have been well documented in the FC literature and should not be entirely surprising given the high degree of individual variability in neuroanatomy and functional networks (Braga et al., 2017. Buckner et al., 2019). Additionally, the QPP algorithm converges on an average pattern no matter what time series it runs on, so it should be no surprise that the group level QPP and underlying network dynamics are not fully recapitulated when running the QPP on a subject-wise level. At the group level, the QPP algorithm was run on the concatenated time series for all 16 subjects from a group at once (14,992 timepoints at once). This means the group QPP converged on an averaged pattern for all 14,992 timepoints for each group. The individual scans’ QPP (937 timepoints per scan) resulted in a distribution of correlations largely consistent with the group level analysis (Figure 3A). However, the group level algorithm may converge on altogether different results depending on the underlying subjects, and because there is much less data in an individual image series. When considering group vs subject-wise QPP results it is also important to note that global signal regressed data has been shown to produce QPPs that may be more similar to one another in terms of individual DMN-TPN correlations (Yousefi et al, 2018).

In summary, the group level QPP correlations we conducted are largely consistent with the seed-based analysis originally done by Belden et al. That is, at the group level, we see the same increase in functional connectivity between the default network and both the visual and frontoparietal networks that were detected by Belden et al. Additionally, an increase in Amyg-DAN connectivity throughout the QPP was found here that was not reported in Belden et al.

### 4.4 Limitations

Small sample size is a common limitation in resting state and task based fMRI interpretation (Thirion et al., 2007. Button et al., 2013. Turner et al., 2018). Given the group sample sizes of n = 16 for this study, it is difficult to draw strong general conclusions regarding creativity. With this type of niche study however, sample size will likely continue to hamper interpretation for the foreseeable future as it is difficult to recruit a large number of age-matched volunteers for a variable such as musical or creative training. Even within this study there is significant heterogeneity of creative training within each group. For example, not all of the improv musicians have the same primary instrument, or the same number or training hours, or the same age of onset for musical training. Attempting to obtain an even larger sample size of musicians would introduce even more variation in the subjects’ respective training backgrounds. However, while small sample size may hinder generalization, in our case it may be beneficial as it shows that our QPP and cPCA methods are robust in a very typical fMRI dataset in terms of sample size.

While the groups used in this study all had the same ratio of male and female subjects (12 and 4, respectively), it would be ideal to have an equal number of male and female subjects overall, as sex differences in rsfMRI have been widely reported (Hjelmervik et al., 2014).

Our primary dynamic FC correlation values are presented on data that were processed with global signal regression (GSR). There is still no consensus on global signal regression as a pre-processing step but the use of global regression has been argued to increase the possibility of detecting spurious anti-correlations (Murphy et al., 2017. Godwin et al., 2017). At the same time, GSR may reveal valid insights into FC patterns missed without GSR (Murphy et al., 2017).

Finally, as noted previously, the primary group level findings were not recapitulated when the QPP algorithm was run on a subject-wise basis. While this is not entirely surprising, the discrepancy between group and individual results shows that we require further understanding of individual FC dynamics. It is possible that individual noise or anatomical variation may very well be greater than subtle but real shifts in network dynamics due to creative training.

### 4.5 Conclusions

This study represents the first simultaneous application of algorithm based and cPCA based QPP analysis to detect group-wise differences in rsfMRI network dynamics. Using cPCA to detect QPPs yielded convergent group level results to those of algorithm-based QPP detection. Improvisational music training was found to be associated with increased visual network and DMN connectivity during quasi-periodic infraslow network dynamics. This is consistent with Belden et al.’s previous static functional connectivity analysis results and extends upon their findings. Both QPP analysis methods support a potential relationship between visual network-DMN connectivity and human creativity. The two methods also found that improvisational music training is associated with higher amygdala and DAN connectivity, further implicating the amygdala in creative cognition.

### 4.6 Future Directions

While the subjects’ age of onset of musical training was available for each subject, the present data are purely cross-sectional. Performing a longitudinal study with two or more scans over the course of musical training would help clarify whether there is a causal link between creative training and altered network dynamics by comparing dynamic DMN FC at different phases of learning. Additionally, comparing the dynamics of artists, writers, or other creatively trained individuals with musical improvisers would help clarify whether the effects of creative training are domain general or differ depending on the creative discipline.

The trend of decreased QPP amplitude in improvisational musicians seems counter to our initial hypothesis that improv musicians may exhibit increased DMN activity. However, the squared difference between DMN and DAN only represents the amplitude of those networks in the infraslow frequency range analyzed in the present study. To better understand the findings of amplitude change in improv musicians, future studies could apply additional bandpasses for internetwork analysis, or include individual comparison of all ROIs within each respective network to provide a sense of which ROIs show altered amplitude during infraslow QPPs or patterns of other frequency ranges.

Because increased amygdala-cortical connectivity is related to anxiety (He et al., 2015), investigating a correlation between trait anxiety, creative training, and dynamic amygdala activity is another opportunity for future studies of dynamic FC and creativity. Creativity and anxiety have been tentatively linked in the literature for some time (Carlsson, 2002. Daker et al., 2020. Vartanian et al., 2020). Such a study of dynamics could be guided using the three amygdala subdivisions established in Sylvester et al., 2020, that is, the default mode, dorsal attention, and unspecified amygdala, to elucidate their respective dynamic FC with cortical networks implicated in creativity.

## Data and Code Availability

(mandatory unless there is no data or code used)

QPP algorithm scripts are available at the Keilholz MIND Lab github here: https://github.com/BnzYsf/QPP_Scripts_v0620

cPCA scripts provided by Taylor Bolt (Bolt et al., 2022) are available at his github here: https://github.com/tsb46/complex_pca

Structural and rsfMRI scans were provided by Alex Belden from the MIND lab at Northeastern and can be made available upon request to the authors and original data collectors.

## Author Contributions

Conceptualization, data preparation, data processing, formal analysis, figure drafting, paper writing, review, editing: HW and AF. Data visualization, analysis, review, and editing: LD. Data preparation and curation: AB, PL. Supervision, review, and editing: SK. Preprocessing troubleshooting, review, and editing: TJL.

## Ethics

All original scan acquisition adhered to best practices for ethical human data collection and informed consent in accordance with local institutional review boards. See original data acquisition study for details (Belden et al., 2020).

## Acknowledgements

We thank the MIND Lab (https://web.northeastern.edu/mindlab/people/) at Northeastern University, Dr. Loui Psyche, Dr. Alex Belden for the collaboration and for the effort in making their musician dataset available to us. We also want to acknowledge the labor of Dr. Behnaz Yousefi who originally worked on updating the QPP code that was eventually modified and used in the present study. We thank Dr. Taylor Bolt for his work on complex PCA and QPPs and his excellent instruction and help regarding the use of his cPCA code for QPP detection.

