## Supplemental Figs 1 and 2 for "Creative tempo: Spatiotemporal dynamics of the default mode network in improvisational musicians"

**Supplementary figures**:

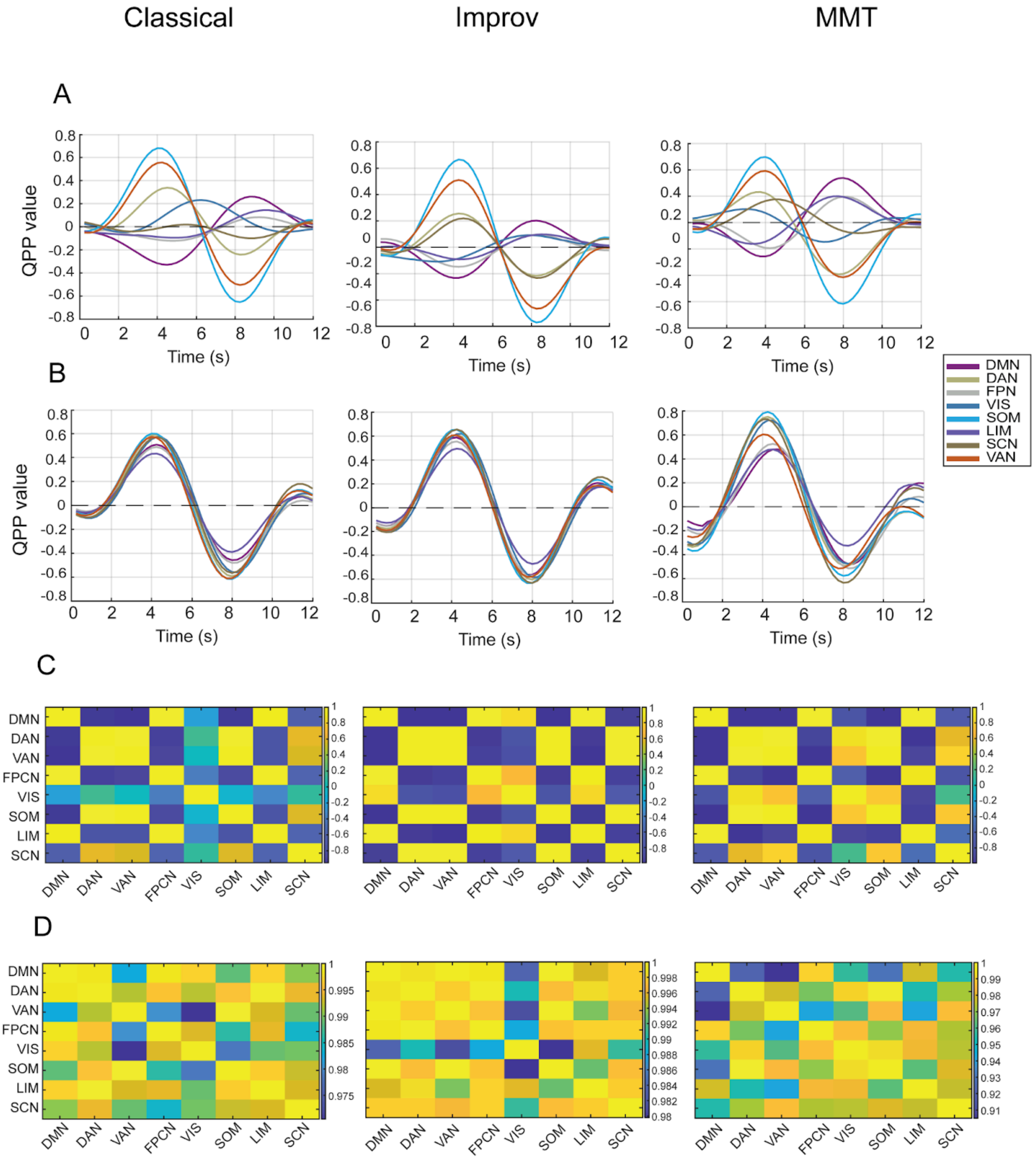


**Supp. Figure 1**: Global signal regressed (GSR) vs non-GSR data. QPP waveforms with GSR (A) and without GSR (B) for the 3 musical training groups. FC matrices showing correlations between 8x8 Yeo's networks during the QPP for GSR (C), and non-GSR (D) scans.

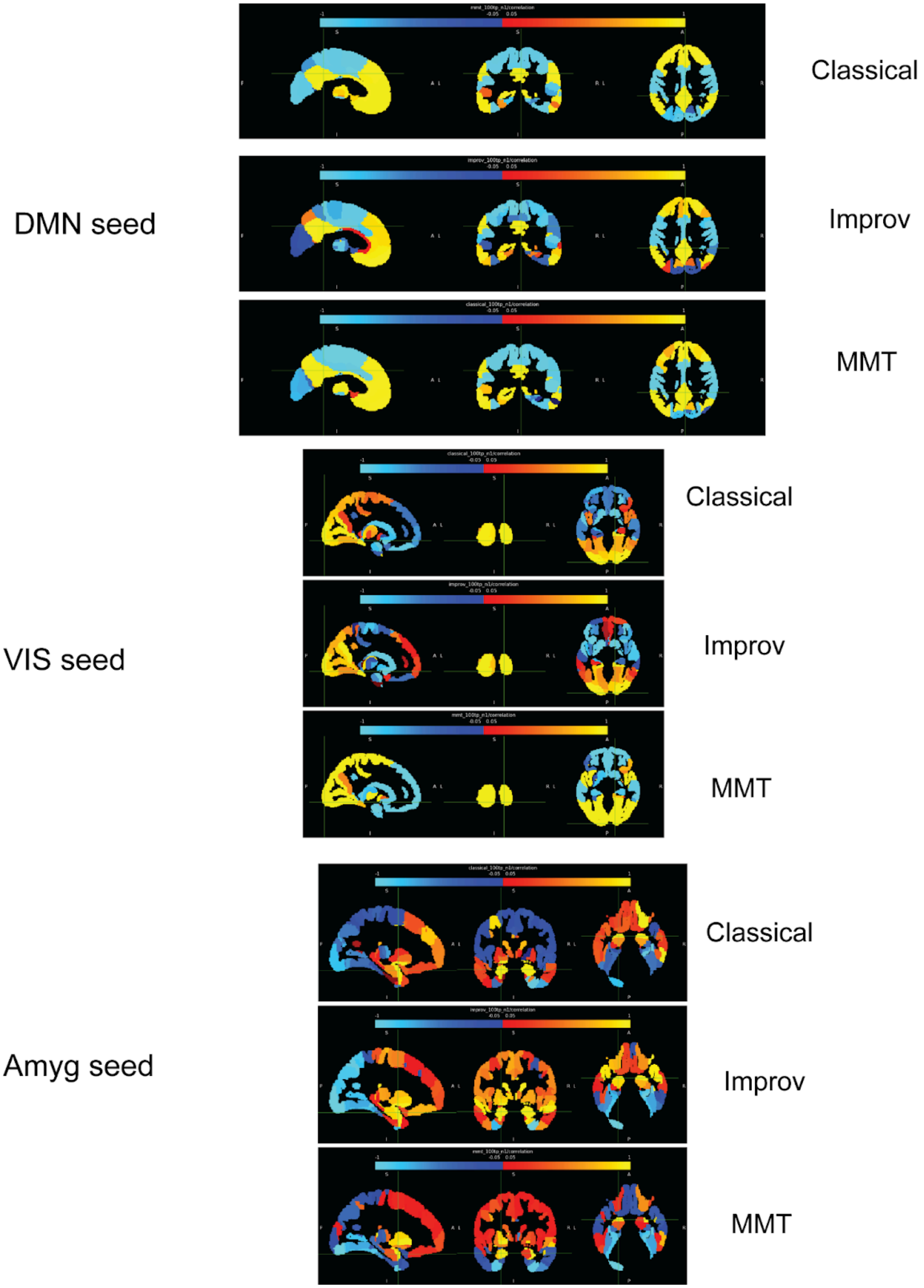


**Supp. Figure 2**: Seed voxel locations within major ROIs analyzed for A) DMN, B), VIS, and C) Amygdala. Color bar indicates correlation to seed region of normalized BOLD signal during 1 cycle of cPCA based QPP between -1 and +1. Note the very high correlation (bright yellow) of all voxels within each ROI to the seed voxels. See table 1 for seed voxel coordinates.
